# An aging-independent replicative lifespan in a symmetrically dividing eukaryote

**DOI:** 10.1101/064832

**Authors:** Eric C. Spivey, Stephen K. Jones, James R. Rybarski, Fatema A. Saifuddin, Ilya J. Finkelstein

## Abstract

The replicative lifespan (RLS) of a cell—defined as the number of generations a cell divides before death—has informed our understanding of the molecular mechanisms of cellular aging. Nearly all RLS studies have been performed on budding yeast and little is known about the mechanisms of aging and longevity in symmetrically dividing eukaryotic cells. Here, we describe a multiplexed fission yeast (*Schizosaccharomyces pombe*) lifespan micro-dissector (FYLM); a microfluidic platform for performing automated micro-dissection and high-content single-cell analysis in well-defined culture conditions. Using the FYLM, we directly observe continuous and robust replication of hundreds of individual fission yeast cells for over seventy-five cell divisions. Surprisingly, cells die without any classic hallmarks of cellular aging such as changes in cell morphology, increased doubling time, or reduced sibling health. Genetic perturbations and longevity-enhancing drugs can further extend the replicative lifespan (RLS) via an aging-independent mechanism. We conclude that despite occasional sudden death of individual cells, fission yeast does not age. These results highlight that cellular aging and replicative lifespan can be uncoupled in a eukaryotic cell.

## Introduction

Aging is the progressive decrease of an organism’s fitness over time. The asymmetric segregation of a set of molecules called pro-aging factors during mitosis has been proposed to promote aging in both yeast and in higher eukaryotes (1–4). In budding yeast, asymmetric division into mother and daughter cells ensures that a mother cell produces a limited number of daughters over its replicative lifespan (RLS) (5). Aging mother cells increase in size, divide progressively more slowly, and produce shorter-lived daughters (5–7). Mother cell decline is associated with asymmetric phenotypes such as preferential retention of protein aggregates, dysregulation of vacuole acidity, and genomic instability (3, 8, 9). By sequestering pro-aging factors in the mothers, newly born daughters reset their RLS (3, 8, 10). These observations raise the possibility that alternate mechanisms may be active in symmetrically dividing eukaryotic cells.

The fission yeast *Schizosaccharomyces pombe* is an excellent model system for investigating RLS and aging phenotypes in symmetrically dividing eukaryotic cells. Fission yeast cells are cylindrical, grow by linear extension, and divide via medial fission. After cell division, the two sibling cells each inherit one pre-existing cell tip (old-pole). The new tip is formed at the site of septation (new-pole). Immediately after division, new growth is localized at the old-pole end of the cell. Activation of growth at the new-pole cell tip occurs ~30% through the cell cycle (generally halfway through G2). The transition from monopolar to bipolar growth is known as new end take-off (NETO; (11–13). Prior studies of fission yeast have yielded conflicting results regarding aging. Early papers reported aging phenotypes akin to those observed in budding yeast (e.g., mother cells become larger, divide more slowly, and have less healthy offspring as they age) (2, 14). However, a recent report used colony lineage analysis to conclude that protein aggregates are not asymmetric distributed, and that inheriting the old cell pole or the old spindle pole body during cell division does not lead to a decline in cell health (15). The apparent controversy between these studies may stem from the difficulty in tracking visually identical cells for dozens of generations. Furthermore, recent work has shown that large sample sizes are needed to truly capture cellular lifespan accurately – populations less than ~100 cells do not reliably estimate the RLS (16).

Here, we report the first high-throughput characterization of both RLS and aging in fission yeast. To enable these studies, we developed the multiplexed fission yeast lifespan microdissector (FYLM) — a high-throughput microfluidic platform and software analysis suite that captures and tracks individual *S. pombe* cells throughout their RLS. Using the FYLM, we observed that the RLS of fission yeast is substantially longer than previously reported (2, 14). Remarkably, cell death is stochastic and does not exhibit the classic hallmarks of cellular aging. Despite the lack of aging phenotypes, rapamycin and Sir2p overexpression both extend RLS, whereas increased genome instability decreases RLS. We conclude that fission yeast dies primarily via a stochastic, age-independent mechanism.

## Results

### A high-throughput microfluidic assay for measuring the replicative lifespan in fission yeast

Replicative lifespan assays require the separation of cells after every division. This is traditionally done via manual micro-dissection of sibling cells, a laborious process that is especially difficult for symmetrically dividing fission yeast. We recently addressed this challenge by developing a microfluidic platform for capturing and immobilizing individual fission yeast cells via their old cell tips (17). However, our first-generation device could only observe a single strain per experiment, required an unconventional fabrication strategy, and suffered from frequent cell loss that ultimately shortened the observation time. To address these limitations, we developed a multi-channel FYLM (multFYLM, **Fig. 1**), along with FYLM Critic—a dedicated software package designed to streamline the analysis and quantification of raw microscopy data. Single cells are geometrically constrained within catch channels, preserving the orientation of the cell poles over multiple generations (**Fig. 1*A, B***). The cells divide by medial fission, thereby ensuring that the oldest cell pole is retained deep within the catch channel. If new-pole tips are loaded initially, then these outward-facing tips become the old-pole tips after the first division. The multFYLM consists of six closely spaced, independent microfluidic subsystems, with a total capacity of 2,352 cells (**Fig. 1*C*, Fig. S1**). To ensure equal flow rates throughout the device, each of the six subsystems is designed to have the same fluidic resistance. Each subsystem consists of eight parallel rows of 49 cell catch channels (**Fig. 1*D***). The catch channel dimensions were optimized for loading and retaining wild type fission yeast cells (**Fig. 1E, Fig. S2;** (17). The eight rows of cell catch channels are arranged between a large central trench (40 μm W × 1 mm L × 20 μm H) and a smaller side trench (20 μm W × 1 mm L × 20 μm H). Cells are drawn into the catch channels by flow from the central trench to the side trenches via a small (2 μm W × 5 μm L × ~12 μm H) drain channel (**Fig. 1F**). We did not monitor all 2,352 catch channels during our experiments, but we routinely filled >80% of the 224 monitored catch channels in each of the six microfluidic sub-systems (**Fig. S2*A***). A constant flow of fresh media supplies nutrients, removes waste, and ensures that cells are stably retained for the duration of the experiment. Once loaded, up to 98% of the cells could be retained in their respective catch channels for over 100 hours (**Fig. S2*B* & *C*, Movie S1**). As a cell grows and divides, the old-pole cell is maintained by flow at the end of the catch channel, while the new-pole cells grow towards the central trench. Eventually, the new-pole cells are pushed into the central trench, where they are washed out by constant media flow (**Movie S1**). This allows for continuous, whole-lifetime observation of the old-pole cell as well as its newest siblings (**Fig. 1*G* & *1H***)

**Fig. 1.**
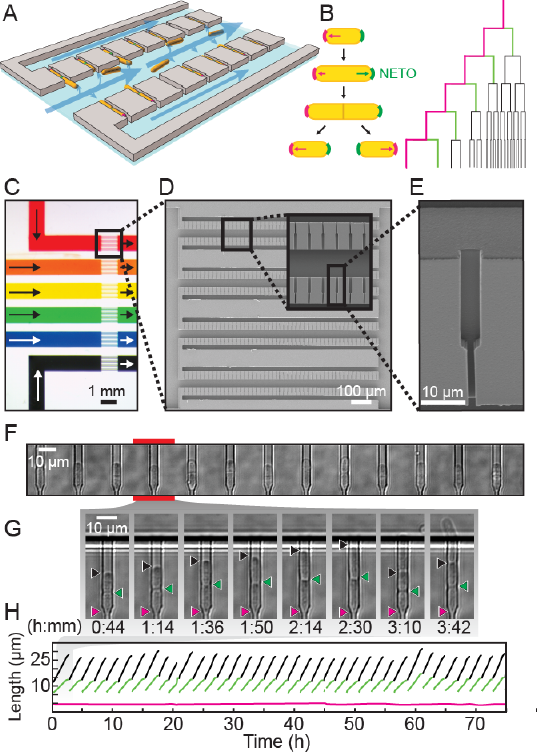
A multiplexed fission yeast lifespan microdissector (FYLM). (A) Illustration of FYLM (gray) loaded with fission yeast (orange). Blue arrows represent media flow though the FYLM. (B) Left: Fission yeast cells initially grow from the old pole end (magenta). After new end takeoff (NETO), growth begins at the new pole end (green). Right: FYLM permits tracking of the old pole cell, as well as its most recent siblings. (C) Multiplexed FYLM (MultFYLM) showing six independent channels. Arrows indicate direction of media flow. Scanning electron micrographs of (D) a single FYLM section and (E) a single catch channel. The channel is long enough to accommodate the old pole cell, as well as the most recent new pole sibling. (F) White-light microscope image of a row of catch channels loaded with cells. (G) Time-lapse images (H) and single-cell traces of a replicating cell. The old pole (magenta) is held in place while the new pole (green) is free to grow. The old pole of the most recent sibling (black) extends until it is removed by flow into the central trench (after ~2 h 30 m).

An inverted fluorescence microscope with a programmable motorized stage allowed the acquisition of single cell data with high-spatiotemporal resolution (**Fig. S3A**). Following data collection, images are processed through the FYLM Critic pipeline (see Supplemental Methods), which measures length and fluorescence information for each cell at each time point (**Fig. S3B**). These data are parsed to determine the cell’s size, division times, growth rates, lifespan, and fluorescence information (if any). Cells within the FYLM grew with kinetics and morphology similar to cells in liquid culture and showed growth and NETO rates (**Fig. S4, Table S1**) similar to those reported previously (15, 18, 19). In sum, the multFYLM allows high content, continuous observation of individual fission yeast cells over their entire lifespans.

### The fission yeast replicative lifespan is not affected by aging

We used the multFYLM to measure the fission yeast RLS (**Fig. 2**). From these data, we plotted the replicative survival curve (**Fig. *2A***) and determined the survival function, *S*(*g*) which is the probability of being alive after generation *g*. Using the survival data, we also computed the hazard function, 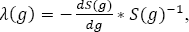, which is the instantaneous risk of death after cell division *g* (**Fig. 2B**). Surprisingly, the fission yeast survival curve did not fit the traditional aging-dependent Gompertz model, (20–22), which describes the RLS in terms of a generation-dependent (*i.e.*, aging) and a generation-independent term (Eq. 2 in Methods). The RLS data was best described by a single exponential decay, corresponding to a generation-independent hazard rate (**Fig. 2B**). Strikingly, the hazard rate does not increase as the replicative age increases; instead, it remains steady at an average 2% chance of death per cell per generation (**Fig. 2*B***) For comparison, we also analyzed the survival data and hazard function for budding yeast (*S. cerevisiae*) grown in a microfluidic device (23). As expected, the budding yeast hazard function increases with each generation and fits the aging-dependent Gompertz model (**Fig. 2**). Thus, the replicative age, *g*, strongly influences the probability of death in budding yeast, but not in fission yeast.

**Fig. 2.**
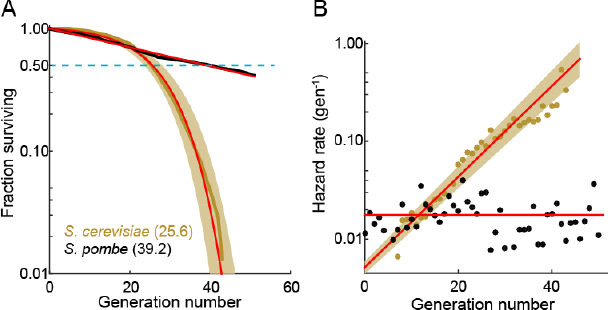
The fission yeast replicative lifespan (RLS). (A) Survival curves for wild-type *S. pombe* (black) and wild-type *S. cerevisiae* (brown, data from Jo et al. 2015); both were grown in microfluidic microdissection devices. Numbers indicate the average lifespan. Red lines are a fit to a Gompertz (*S. cerevisiae*) and exponential decay (*S. pombe*) survival models. Shading indicates 95% confidence interval (C.I.). Dashed blue line: 50% survival. (B) Hazard rate curves for the data shown in (A). The hazard rate increases dramatically with increased replicative age for *S. cerevisiae* but not for *S. pombe*.

The RLS is defined by the number of generations at which 50% of the starting cells are dead. We measured an RLS of 39.2 generations (95% C.I. 38.6-39.8, n=440, **Fig. 2A**) for the laboratory strain 972h-grown in rich media (24). To explore the effect of different genetic backgrounds on replicative lifespan, we determined survival curves, hazard functions, and replicative half-lives for three additional strains, including two wild fission yeast isolates with distinct morphologies and doubling times (CBS2777 and JB760; (25). Most laboratory fission yeast strains have three chromosomes, but the recently identified strain CBS2777 has four (26). The fourth chromosome in CBS2777 originated by a series of complex genomic rearrangements (27). CBS2777 had the shortest RLS of 22.7 generations (95% CI 21.9-23.6 generations), consistent with its aberrant genome and altered genome maintenance (27). In contrast, strains NCYC132 and JB760 showed greater longevity with estimated RLS of >50 and >70 generations, respectively (**Fig. S5, Table S2**). These RLS correspond to a hazard rate that is ~5-fold lower than strain h-972 (**Fig. S5B**). The longevity of strain NCYC132 is consistent with an earlier manual microdissection study performed on solid media (15). Each strain’s survival curve was best described by an exponential decay function, resulting in a generation-independent hazard function (**Fig. S5**, **Table S2**). Thus, an aging-independent replicative lifespan is a feature of diverse fission yeasts isolates, and likely the entire species.

### Aging-associated phenotypes do not correlate with death in fission yeast

Prior RLS studies have identified three common phenotypes associated with cellular aging (28, 29). In budding yeast, aging mother cells increase in cell size, progressively slow their doubling times, and produce daughters with decreased fitness (5–7) . Whether older fission yeast cells also undergo similar aging-associated phenotypes remained unresolved. To answer these outstanding questions, we examined time-dependent changes in morphology, doubling times, and sibling health in individual fission yeast cells as they approached death. Quantitative observation of cell length revealed two classes of dying cells: the majority (72%; n=234) died prior to reaching the normal division length (< 16.2 μm, **Fig. 3*A* & 3*B***). The remaining cells (28%; n=92) showed an elongated phenotype and exceeded the average division time by more than three-fold. In *S. pombe*, cell length is strongly correlated with division time and cell length, indicating that cell cycle checkpoints were likely dysregulated in these dying cells (11, 12, 30, 31). Despite these differences in length at death, most cells had normal doubling times throughout their lifespan (**Fig. 3*C* & 3*D***). Next, we investigated whether there were any morphological changes in the generations preceding death. The majority of cells retained wild type cell lengths and doubling rates until the penultimate cell division. In the two generations immediately preceding death, there was a mild, but statistically significant change in the distributions of cell lengths (**Fig. 3*E***) and doubling times (**Fig. 3*F***). However, we did not observe any predictable trends for individual cells, arguing against a consistent pattern of age-related decline. In sum, these results strongly support the conclusion that *S. pombe* dies without aging-associated morphological changes.

**Fig. 3.**
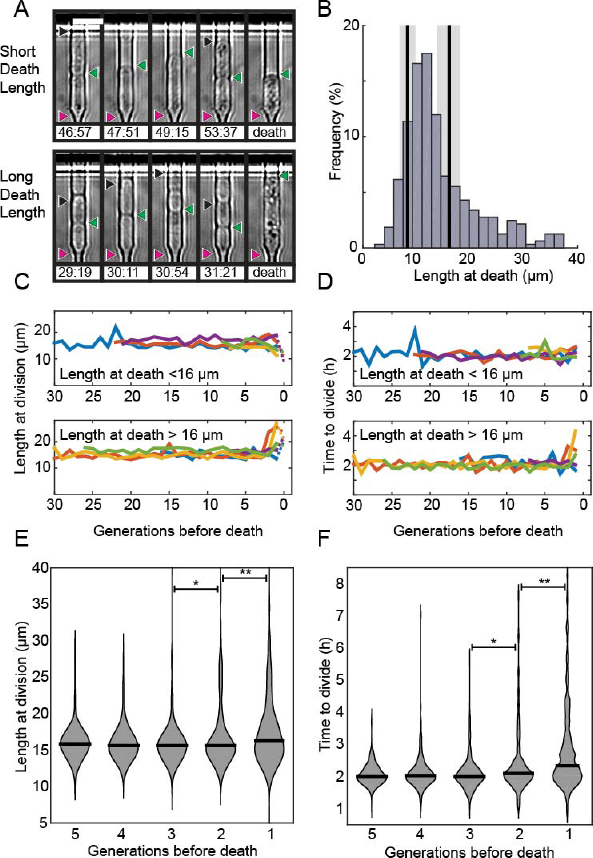
Fission yeast does not show signs of aging. (A) Images of cells showing a short (top) and long (bottom) phenotype at death. Triangles indicate the old-pole, new-pole, and the new-pole of the previous division as in Fig. 1*G*. Scale bar: 10 μm. (B) Histogram of cell length at death. The birth length was 8.3 ± 1.5 μm (mean ± st. dev., n=326) and the length at division was 16 ± 2.2 μm (n=326). (C) Cells dying with either short (top) or long (bottom) phenotype have normal length and (D) doubling times prior to death, as indicated by five representative cells. (E) Distribution of length and (F) doubling time at division in the five generations preceding death. Cells were post-synchronized to the time of death. The black bar shows the median value for each generation (n > 290 cells for all conditions). Sequential generations were compared using the Kolmogorov-Smirnov test (* for p<0.05, ** for p<0.01).

We next determined the fate of the last sibling produced by dying cells. This is possible because the last siblings of a dying cell also remain captured in the catch channels (**Fig. 4A**). These are siblings that are produced by the final division of the old-pole cell. We categorized these siblings into three classes: (i) those that died without dividing, (ii) those that divided once and died, and (iii) those that divided two or more times (**Fig. 4*A*, Movies S2-S4**). The fate of the new-pole siblings was independent of the replicative age of the old-pole cells at death (**Fig. 4*B***). Most new-pole siblings of a dying cell (66%, n=135) never divided and typically did not grow, suggesting that the underlying cause of death was distributed symmetrically between the two sibling cells. Similarly, new-pole siblings that divided once and died (14%, n=29) also typically did not grow. Old pole cells that died while hyper-elongated were more likely to have these unhealthy offspring (**Fig. 4*C*, 4*D***). As elongated cells generally indicate the activation of a DNA damage checkpoint (32), our findings suggest that genome instability in these cells may be the underlying cause of death in both the old cell and its most recent sibling. In new-pole siblings that divided two or more times (20%, n=40), doubling times were indistinguishable from rapidly growing, healthy cells (2.1 ± 0.7 hours, n=208). These healthy new-pole cells were typically born from old-pole cells that did not elongate during their terminal division (**Fig. 4*C* & 4*D***). These observations suggest that the cause of death was contained within the old-pole cell, thereby reducing the likelihood of death in the sibling. Taken together, our observations suggest that in the majority of cases, cell death impacts both sibling cells. However, in 24% of cells, death is localized to just one of the two siblings. Importantly, there was no correlation between the replicative age of the old-pole cell and survival probability of the last sibling cell. These observations are in stark contrast to *S. cerevisiae*, where aging factors are generally sequestered to younger mother cells (3, 33). However, older *S. cerevisiae* mothers produce larger and slower-dividing daughter cells. Thus, we do not observe any aging-dependent outcomes for the fate of the new-pole sibling, further confirming that *S. pombe* does not age.

**Fig. 4.**
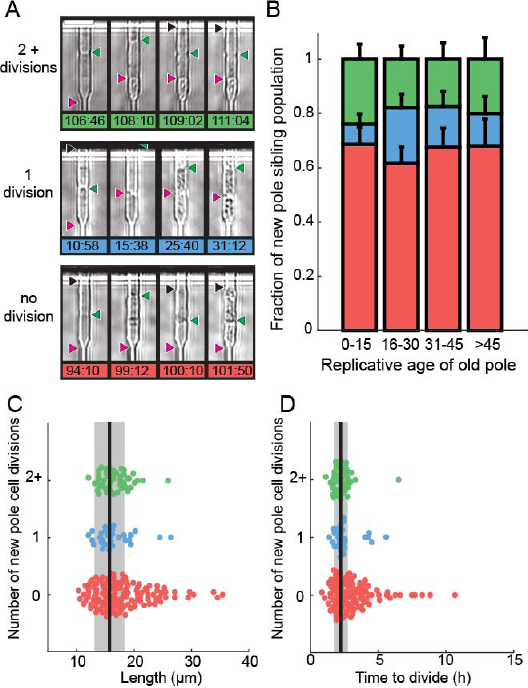
Analysis of siblings born during the last division of a dying cell. (A) Last new-pole sibling continued to divide (top), died after one division (middle) or died without dividing (bottom). (B) The distribution of last-sibling phenotypes as a function of the old pole replicative age (n=245). Error bars are st. dev. measured by bootstrap analysis. (C) The length at division and (D) the doubling time of the new-pole siblings. Vertical black lines and gray bars show the mean and standard deviation of the total cell population.

### Genetic manipulation and rapamycin treatment extend replicative lifespan

Although *S. pombe* does not show aging phenotypes, the replicative lifespan may still be extended via chemical or genetic manipulations. The histone deacetylase Sir2p has been shown to modulate lifespan and aging in a wide variety of organisms from yeasts to mice (34–36). For example, *sir2* deletion (*sir2Δ*) in budding yeast reduces the replicative lifespan by ~50% (23, 37), whereas Sir2p overexpression can increase the RLS by up to 30% (37). The *S. pombe* genome encodes three Sir2p homologs, one of which shares a high degree of sequence similarity and biochemical functions with the budding yeast Sir2p (2, 38). We constructed strains of fission yeast with deletion and overexpression of this most similar Sir2p homolog. The RLS of the *sir2Δ* strain was 33.4 generations (95% CI 32.6-34.2, n=329), which was comparable to that of the wild type strain (39.2 generations, 95% CI 38.6-39.8, n=440; **Fig. 5*A***). These results are consistent with prior observations that fission yeast wild type and *sir2Δ* cells had similar growth rates when cultured without stressors (2, 39). In contrast, constitutive 6-fold *sir2* overexpression (*sir2OE*, **Fig. S6)** increased the RLS over 50%, with a mean replicative lifespan of >60 generations (n=301; **Fig. 5*A***). Cells overexpressing Sir2p have a 2-fold lower hazard rate (total risk of death) than wild-type cells **(Figs. 5*B* and S7**). In sum, overexpression of Sir2p increases the RLS of *S. pombe*, but does not affect aging.

**Fig. 5.**
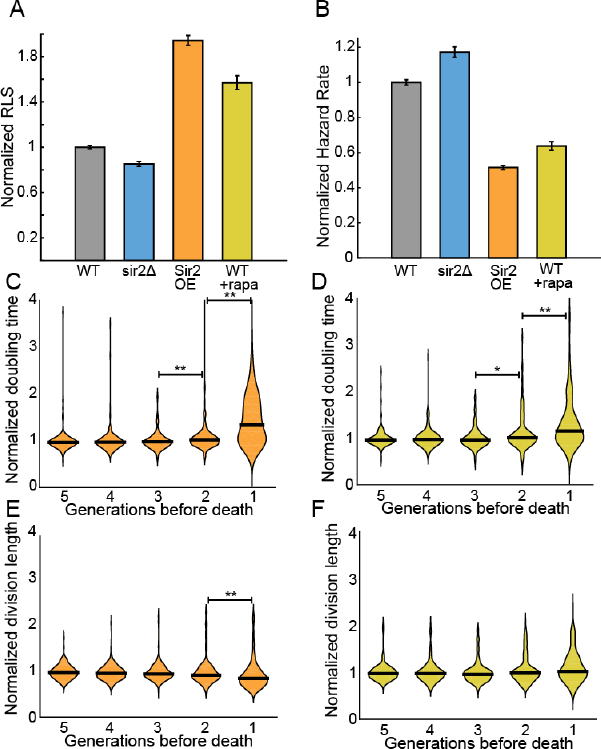
Sir2p and rapamycin extend replicative lifespan. (A) Replicative lifespans and (B) hazard rates of strains normalized to the mean RLS and hazard rate of wild-type (h-972) strain. Error bars: 95% C.I. on an exponential decay fit to the experimental survival curve. (C) Distribution of normalized doubling time in the five generations preceding death for Sir2p overexpression cells and (D) wild-type cells treated with 100 nM rapamycin. (E) Distribution of normalized length at division in the five generations preceding death for Sir2p overexpression cells and (F) wild-type cells treated with 100 nM rapamycin. Black bars show the median value. Sequential generations were compared using the Kolmogorov-Smirnov test (* for p<0.05, ** for p<0.01; n>144 cells for all conditions).

Rapamycin inhibits target of rapamycin (TOR) proteins in eukaryotes (40, 41), which are key modulators of aging and age-related diseases across many model systems (42). Rapamycin increases the replicative lifespan of budding yeast (43), and also increases the longevity of worms, flies and mice (44–46). However, the effect of rapamycin on fission yeast RLS has not been reported. Addition of 100 nM rapamycin to the flow medium increased the RLS by 41% (55.3 generations, 95% CI 53.1-57.6, n=184; **Fig. 5*A***). As with Sir2p overexpression, treating cells with rapamycin increased their longevity by reducing the aging-independent hazard rate (**Fig. 5*B***). Importantly, both rapamycin and Sir2p overexpression did not affect the lengths or doubling times of cells as they approached death. The cell length at division remained constant for the five generations preceding cell death, whereas the cell doubling time increased only in the last generation before death **(Fig. 5*C*-*F***). These results are consistent with an aging-independent mechanism for RLS extension in *S. pombe*.

### Ribosomal DNA (rDNA) instability contributes to sudden cell death

Ribosomal DNA (rDNA) is a repetitive, recombination-prone region in eukaryotic genomes. Defects in rDNA maintenance arise from illegitimate repair of the rDNA locus and have been proposed to be key drivers of cellular aging (36, 47). Fission yeast encodes ~117 rDNA repeats on the third chromosome, and incomplete rDNA segregation can activate the spindle checkpoint (48, 49). If unresolved, this can result in chromosome fragmentation and genomic instability (50). Indeed, we observed that 28% of dying cells were abnormally long, suggesting activation of a DNA-damage checkpoint (**Fig. 3B**). Aberrant rDNA structures are processed by the RecQ helicase Rqh1p and mutations in the human Rqh1p homologs are implicated in cancer and premature aging-associated disorders (51–55). Therefore, we sought to determine whether rDNA instability and loss of Rqh1p can contribute to stochastic death.

We visualized segregation of rDNA in cells using the nucleolar protein Gar2 fused to mCherry. Gar2 localizes to rDNA during transcription and has been used to monitor rDNA dynamics in live cells (50). The strain expressing Gar2-mCherry at the native locus divided at approximately the same length and rate as wild-type cells (**Fig. S8**). As expected, cells mostly had equal Gar2 fluorescence in both sibling cell nuclei (**Figs. 6*A*-*B***). However, 7% of dividing cells (n=88/1182) showed rDNA segregation defects that appear as multiple rDNA loci, asymmetric fluorescence distributions, or rDNA bridges (**Fig. 6*A***). Multi-punctate rDNA loci were nearly always lethal, whereas asymmetric rDNA segregation and rDNA bridges were lethal in 40% of cells (n=31/78; **Fig. 6*B***). As expected, *rqh1Δ* cells showed higher frequencies of spontaneous rDNA defects, a shorter RLS (5.3 generations, 95% CI 5.0-5.7), and a > 7-fold higher hazard rate (**Fig. S9**). Remarkably, rDNA defects were highly elevated in wild type cells immediately preceding death (**Fig. 6C**), and were even more prevalent in dying *rqh1Δ* cells (**Fig. S9**). Thus, Rqh1p promotes the longevity of fission yeast by suppressing rDNA instability.

**Fig. 6.**
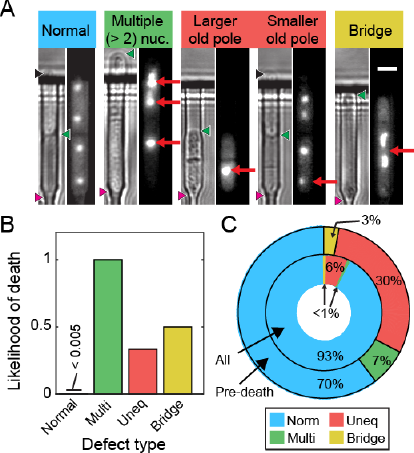
Ribosomal DNA (rDNA) instability is highly correlated with cell death. (A) Images of cells exhibiting rDNA instability, as reported by gar2-mCherry, which binds to rDNA. (B) Likelihood of cell death following one of the defects observed in (A). (C) Dying cells exhibited elevated rDNA defects (outer ring) relative to healthy dividing cells (inner ring).

## Discussion

Here, we report the first study of fission yeast RLS in a high-throughput microfluidic device (**Fig. 1**). The multFYLM provides a temperature-controlled growth environment for hundreds of individual cells for up to a week (>75 generations), facilitating tens of thousands of micro-dissections. We also developed FYLM Critic, an image-processing pipeline for quantitative phenotypic analysis of individual cell lineages. We note that microfluidics-based lifespan measurements are free from potential extrinsic effects (e.g., secretion of small molecules onto the solid agar surface) which have been proposed to confound observations of RLS assays using manual microdissection (56). Using the multFYLM, we set out to determine whether fission yeast undergoes replicative aging.

Taken together, our data provide three sets of experimental evidence that fission yeast does not age. First, the RLS survival curves and corresponding hazard rates are qualitatively different between budding and fission yeasts (**Fig. 2**). The budding yeast RLS is best described by a Gompertz model that includes both an age-dependent as well as an age-independent survival probability. In contrast, the fission yeast RLS is best described by a single exponential decay. This corresponds to a hazard function *h(g)* that is constant with age, as would be expected for an organism that does not age. Second, we observed that cell volumes and division rates did not change until the penultimate cell division (**Fig. 3**). Third, after the death of a cell, the health of the surviving sibling was also independent of age (**Fig. 4**). These traits were observed in both laboratory-derived and wild-type fission yeast isolates. (**Fig. S5**). Our conclusions are also broadly consistent with a recent report that observed the phenotypic changes in dying fission yeast cells grown in microcolonies on solid media (15).

Our results highlight a critical consideration when interpreting replicative lifespan curves. The experimentally determined RLS does not necessarily report on whether a population of cells is aging. This is because the RLS is affected by a combination of both age-independent and age-dependent biological processes—cell may die randomly without aging. A mathematical model is required to further parse the relative contributions of aging-independent and aging-dependent mechanisms. The Gompertz function is an excellent model for understanding the replicative lifespan survival curve (57). In this model, the replicative lifespan is dependent on just two parameters: (i) an age-independent hazard rate, and (ii) an age-dependent hazard rate, which determines how rapidly mortality increases as a function of the cell’s age (58). Together, these two parameters describe a population’s likelihood of death at any age. Figure 7 illustrates that the experimentally observed RLS is dependent on both of these terms. An increased RLS is not on its own sufficient to conclude that cells age more slowly. Indeed, our data indicate that a decreased RLS can also be achieved by increasing the age-independent hazard rate. To further illustrate this point, we include a Gompertz model analysis of the RLS *of S. cerevisiae* grown in a microfluidic device (23) and our own data from *S. pombe*. For all of our data, changes in RLS can be completely accounted for by changes in the age-independent hazard rate. In contrast, budding yeast have age-dependent hazard rates an order of magnitude higher than their age-independent rates (**Fig. 7**). We hope that the development of the multFYLM, as well as a quantitative analysis framework, will continue to inform comparative RLS studies in model organisms.

**Fig. 7.**
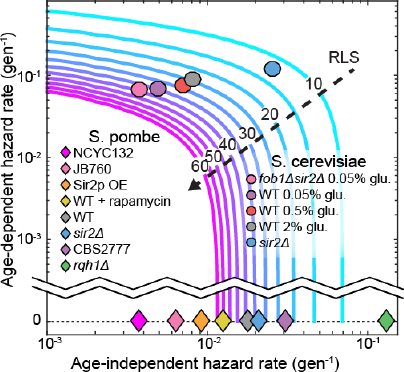
The replicative lifespan is an incomplete reporter of cellular aging. RLS contours were generated from experimentally determined ranges of Gompertz coefficients using Equation [7]. Fission yeast strains examined in this study and budding yeast from an analogous study (23) were plotted on the chart based on the coefficient values from either a Gompertz or exponential decay fit. In all cases, the 95% CI of the coefficients was smaller than the marker size.

What are the molecular mechanisms of stochastic cell death in a symmetrically dividing eukaryote? Strikingly, the RLS of *S. pombe* can be extended by up to 41% via rapamycin treatment and Sir2p over-expression without influencing aging-related phenotypes (**Fig. 5**). This suggests that rapamycin and Sir2p both reduce the onset of sudden death. For a subset of cells, genomic instability due to aberrant segregation of the rDNA locus may be one cause of death (9, 36). Consistent with this hypothesis, rDNA segregation defects were highly elevated in unstressed wild type cells immediately prior to death (**Fig. 6**). Ablating Rqh1p, a helicase that promotes rDNA stability and replication fork progression (59–61), further increased rDNA defects while drastically reducing the RLS, independent of age. Recent studies also indicate that protein aggregation may be a second major source for sudden death in rapidly dividing fission yeast cells (15, 62). Additional studies will be required to determine whether protein aggregates can trigger aging-dependent phenotypes in fission yeast. The mechanisms of death in fission yeast, be they driven by genome instability, protein aggregation or other factors, need not be mutually exclusive. Further study will ultimately define the molecular mechanisms of aging-independent death in fission yeast.

## Materials

### FYLM master structure photolithography

Master structures were fabricated using SU-8 3005 photoresist for the first layer and SU-8 2010 photoresist for the second layer, following standard photolithography techniques described in the product literature (Microchem, Westborough, MA). Photoresist was spun onto 100 mm-diameter, test-grade, P-doped silicon wafers (ID# 452, University Wafers). Photoresist thickness was 20-30 μm for the first layer (conduits, central and side trenches), and 5-6 μm for the second layer (catch and drain channels). Custom chrome on quartz photomasks (Compugraphics) were manufactured from designs created using the freeware integrated circuit layout editor Glade (www.peardrop.co.uk) or with OpenSCAD (www.openscad.org). A Suss MA-6 Mask Aligner (Suss MicroTec Lithography GmbH) was used for mask alignment and photoresist exposure.

### FYLM construction

Approximately 25 g of polydimethylsiloxane (PDMS, Sylgard 184, Dow Corning) was mixed at a weight ratio of 10:1 polymer:hardener, placed on a rotator for >30 minutes, then centrifuged to remove bubbles. A tape barrier was applied to the edge of a silicon wafer bearing the FYLM master structures, and 13 g of PDMS was poured onto the wafer, covering the surface. The wafer with PDMS was degassed for ~10 minutes (at 60-70 cmHg vacuum in a vacuum chamber) to remove additional bubbles, then placed in a 70°C oven for 15-17 minutes. The wafer was intentionally removed while the PDMS was still tacky to improve adhesion of the microfluidic connectors (nanoports). To create a source and drain interface for the device, nanoports (N-333-01, IDEX Health and Science, Oak Harbor, WA) were applied to the surface of the PDMS over the ends of the master structure of the FYLM, then an additional 14 g of PDMS was poured over the surface, with care taken to avoid getting liquid PDMS inside the nanoports. The wafer with PDMS and nanoports was then degassed >10 minutes to remove additional bubbles, and then returned to the 70°C oven for 3 hours to cure completely. The cured PDMS was then removed whole from the wafer after cooling to room temperature. A FYLM with nanoports was trimmed from the cured PDMS disk to approximately 15 × 25 mm. A 1 mm diameter biopsy punch (Acupunch, Accuderm) was used to make holes connecting the bottom of each nanoport to the conduits on the bottom surface of the PDMS. The FYLM was then placed in isopropanol and sonicated for 30 minutes, removed, and placed to dry on a 70°C hotplate for 2 hours. Borosilicate glass coverslips (24 × 60 mm #1, Fisher for FYLM, or 48 × 65 mm #1, Gold Seal for multFYLM) were concurrently prepared by cleaning for >1 hour in a 2% detergent solution (Hellmannex II, Hellma Analytics), then rinsing thoroughly in deionized water and isopropanol before drying for >2 hours on a 70°C hotplate. Cleaned FYLM and coverslips were stored in a covered container until needed. To bond, the FYLM and a coverslip were placed in a plasma cleaner (PDC-32G, Harrick Scientific) and cleaned for 20 seconds on the “high” setting with oxygen plasma from ambient air (~21% oxygen). The FYLM and glass coverslip were then gently pressed together to form a permanent bond. To make the microfluidic interface, PFA tubing (1512L, IDEX), with a 1/16” outer diameter was used to connect a 100 mL luer lock syringe (60271, Veterinary Concepts, www.veterinaryconcepts.com) to the FYLM, and to construct a drain line leading from the FYLM to a waste container. The tubing was connected to the FYLM nanoports using 10-32 coned nuts (F-332-01, IDEX) with ferrules (F-142N, IDEX). A two-way valve (P-512, IDEX) was placed on the drain line to allow for better flow control. For longer experiments, two syringes could be loaded in tandem, connected with a Y-connector (P-512, IDEX). The valve and Y-connector were connected to the tubing using ¼-28 connectors (P-235, IDEX) with flangeless ferrules (P-200, IDEX). The source line tubing was connected to the syringe with a Luer adapter (P-658, IDEX). The syringe(s) was/were then placed in a syringe pump (Legato 210, KD Scientific) for operation. A 2 μm inline filter (P-272, IDEX) was placed upstream of the FYLM to reduce the chance of debris clogging the FYLM during operation.

### Loading the FYLM with cells

As soon as possible after plasma bonding (usually within 15 minutes), the FYLM was placed on the microscope stage and secured, then 15 μL of cells suspended in YES media with 2% BSA were gently injected into the source nanoport. The drain and source interface tubing (with drain valve closed) were then connected to the drain and source nanoports. The syringe pump was then started at 40 μL min^−1^, and the drain valve was opened to allow flow to start. The syringe pump was adjusted between 20-60 μL min^−1^ until flow was established, and catch channels were observed to be filling with cells, then a programmed flow cycle was established: 1-5 minutes at 50 μL min^−1^ followed by 10-14 minutes at 5 μL min^−1^ (average flow rate, 0.5-1.2 mL h^−1^).

### Time-lapse imaging

Images were acquired using an inverted microscope (Nikon Eclipse Ti) running NIS Elements and equipped with a 60X, 0.95 NA objective (CFI Plan Apo λ, Nikon) and a programmable, motorized stage (Proscan III, Prior). The microscope was equipped with Nikon’s “Perfect Focus System” (PFS), which uses a feedback loop to allow consistent focus control over the multiple-day experiments. Images were acquired approximately every two minutes using a scientific-grade CMOS camera (Zyla 5.5, Andor). To improve contrast during image analysis, images were acquired both in the plane of focus and 3-4 μm below the plane of focus. Temperature control was maintained using an objective heater (Bioptechs) and a custom-built stage heater (Omega) calibrated to maintain the FYLM at 30-31°C.

Fluorescence images were acquired with a white light LED excitation source (Sola II, Lumencorp) and a red filter set (49004, Chroma). Fluorescent images were acquired concurrently with white light images, but only every four minutes.

### Media and Strains

All strains were propagated in YES liquid media (Sunrise Scientific) or YES agar (2%) plates, except where otherwise noted. A complete list of strains is reported in Supplemental Table 3. Deletion of *sir2* in the wild-type 972h-strain was completed by first generating a PCR product from IF37 containing KANMX4 flanked by *SIR2* 5’ and 3’ untranslated regions. Competent IF30 cells were then transformed with the PCR product and selected on YES+G418 agar plates. Deletion was confirmed by PCR. A *SIR2* over-expression plasmid was generated via Gateway cloning (Life Technologies). First, *SIR2* was integrated into the pDONR221 plasmid to yield pIF133. *SIR2* was then swapped from pIF133 to pDUAL-HFF51c (Riken), yielding plasmid pIF200. pIF200 expresses *SIR2* under the constitutive *TIF51* promoter, and also labels the protein C-terminally with a His6-FLAG-FLAG tag. Plasmid pIF200 was then transformed into strain IF140 for integration at the *leu1* locus (IF230, IF231; 63). Integration was checked by PCR and expression was determined by RT-qPCR.

### FYLM Critic: An open-source image processing and quantification package

To analyze our single-cell data, we developed FYLM Critic—a Python package for rapid and high-content quantification of the time-lapse microscopy data (github.com/finkelsteinlab/fylm). The FYLM Critic pipeline consists of two discrete stages: (i) pre-processing the raw microscope images to correct for stage drift, and (ii) quantification of cell phenotypes (e.g., length, doubling time, fluorescence intensity and spatial distribution).

### Stage 1: Pre-processing the microscope images

Several image pre-processing steps were taken to permit efficient quantification.

First, the angles of the four solid sections of PDMS on either side of the catch channels were measured relative to the vertical axis of the image, and a corrective rotation in the opposite direction was applied to all images in that field of view. This resulted in the long axis of each cell being completely horizontal in all subsequent analyses. The FYLM was too large to be imaged in its entirety within a single field of view, so the microscope stage was programmed to move to eight different subsections of the device in a continuous cycle. Due to the imperfect motion of the microscope stage, subsequent images of the same field of view randomly translated by a few micrometers relative to the previous image at the same nominal location. Images were aligned to the first image of the sequence using a cross-correlation algorithm (64). The exact location of each catch channel was manually specified, but only if a cell had already entered the channel at the beginning of the experiment.

### Stage 2: Annotating individual cells within the FYLM

In order to analyze cell lengths and phenotypes, kymographs were produced for each catch channel and then annotated semi-manually. To produce the kymographs, a line of pixels was taken along the center of each catch channel, starting from the center of the “notch” and extending into the central trench. This kymograph captured information along the long axis of the cells. The time-dependent one-dimensional (1D) kymographs for each catch channel were stacked together vertically, producing a two-dimensional (2D) kymograph of each cell’s growth and division. The out-of-focus image was used for this process as the septa and cell walls were much more distinct, as described previously (19). The position of the old pole and the leftmost septum were then manually annotated using a simple point-and-click interface to identify each cell division. Once all single-cell kymographs were annotated, this annotation was used to calculate a cell length for each time point. The final status of the cell (whether it died, survived to the end of the experiment, or was ejected) was determined by the user. Further analysis of the cell length, growth rates, and fluorescence intensities was performed in MATLAB (Mathworks). The local minima of the growth curves were used to determine division times and the replicative age of each cell.

Ejections and replacement of individual cells within the catch channels may confound our whole-lifespan tracking. Therefore, we optimizing cell loading, retention and imaging conditions to minimize the loss of individual cells throughout the course of each experiment. First, ejection of individual cells was minimized over the course of ~100 hours (**Fig. S2*B***). Second, each cell was imaged once every two minutes, minimizing the amount of time during which a cell could conceivably be ejected and replaced by another from the central trench. If a cell were indeed replaced within our two-minute frame rate, we would expect an abrupt change in the captured cell length (indicated a re-loading event). All kymographs were carefully monitored for such events, and in the rare cases where this occurred, were discarded from the final analysis. In summary, our analysis ensures that cell ejection or re-capture is mitigated by both efficient cell capture and by analysis of the single-cell datasets.

## Survival curves

Kaplan-Meier (65) survival function estimates were calculated in MATLAB. Due to the low number of lost cells (**Fig. S2*B***) and the high total number of total cells (n ≥100 for all experiments), the survival curves did not include cells that were lost during the course of the experiment (i.e., there was no right censoring). Our analysis is consistent with similar studies in *S. cerevisiae* (66). The survival function estimates were fit with either a simple exponential decay function

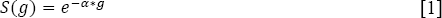

or the Gompertz survival function (20, 21)

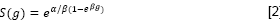

Where S is the fraction of the initial population surviving at generation *g*. The coefficients α & β are >0 and <1, and are further defined for the hazard function λ(g) below. The plotted survival data were weighted at to increase the influence of older cells on the fit.

The hazard function λ(g) (the probability of cell death during a given generation) can be derived from the survival function:

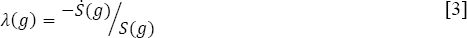

where *S* is the first derivative of *S*. For a survival curve fit to an exponential decay function, the corresponding hazard function is a constant:

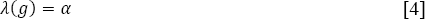

For a survival curve fit to the Gompertz function, the hazard function simplifies to:

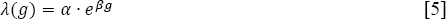

Where the coefficient α amplitude-scales λ(g), and so has an age-independent influence on the hazard function, while the coefficient β time-scales λ(g), and so provides a compact method to describe the age-dependent increase in the probability of cell death (Fig. 7). Note that as *β* → 0, Equation [5] approaches Equation [4].

Mean replicative lifespan (RLS) can be calculated by setting S(g) = ½, and solving for *g* (67). For exponential decay, RLS is simply the half-life of the decay function:

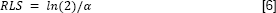

For the Gompertz function, RLS is:

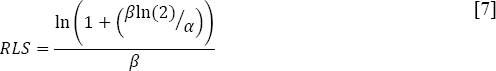

## Statistical Data Analysis

All fitting and statistical data analysis was performed in MATLAB. Briefly, datasets were first tested for normality using the Anderson-Darling method, and parametric tests were used when possible. Fitting of all data was performed in MATLAB using either the nonlinear least squares method or least squares method. Determination of whether a survival curve was fit to an exponential decay or Gompertz function was made primarily by selecting the fit with the highest adjusted r^2^ value.

## Author Contributions

E.C.S., S.K.J., J.R.R. and I.J.F designed research, E.C.S. and S.K.J. performed research, J.R.R. wrote software, E.C.S, S.K.J., J.R.R. and F.A.S. analyzed data; and E.C.S, S.K.J., J.R.R. and I.J.F. wrote the paper.

The authors declare no conflict of interest. This article contains supporting information online.

## Acknowledgments

We are indebted to our colleagues Jürg Bähler, Megan King, Edward Marcotte, Akihisa Matsuyama, Ronit Weisman, and Blerta Xhemalce for valuable strains and reagents. We are grateful to Xenia Brianna Gonzalez for her assistance with data quantification, and to other members of the Finkelstein laboratory for carefully reading the manuscript. We thank our colleagues Edward Marcotte, Andreas Matouschek and Aashiq Kachroo for critical feedback. This work was supported by the American Federation of Aging Research (AFAR-020), the National Institute of Aging (F32 AG053051 to S.K.J.), CPRIT (R1214 to I.J.F.), and the Welch Foundation (F-l808 to I.J.F.). I.J.F. is a CPRIT Scholar in Cancer Research. The content is solely the responsibility of the authors and does not necessarily represent the official views of the National Institutes of Health.

---

**Supplemental Figure 1.**
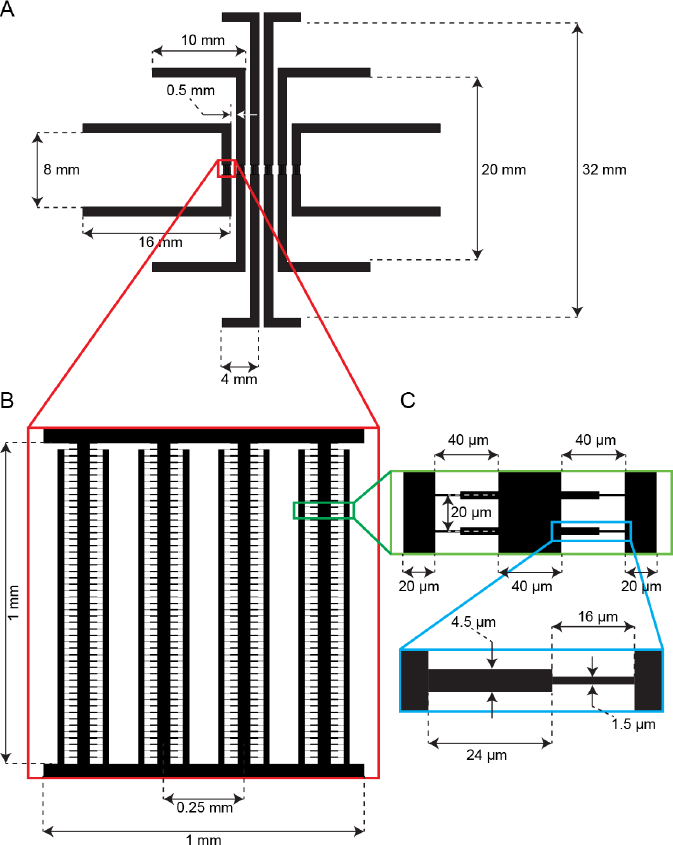
Multiplexed FYLM (MultFYLM). (A) The parallel FYLM subunits are designed to have the same fluidic resistance. This ensures similar flow rates through all devices when equal pressure is applied. (B) Detail showing the arrangement of central and side trenches of a single FYLM subunit. (C) Detail showing arrangement of the catch channels relative to the central and side trenches. (D) Detail showing the dimensions of one catch channel.

**Supplemental Figure 2.**
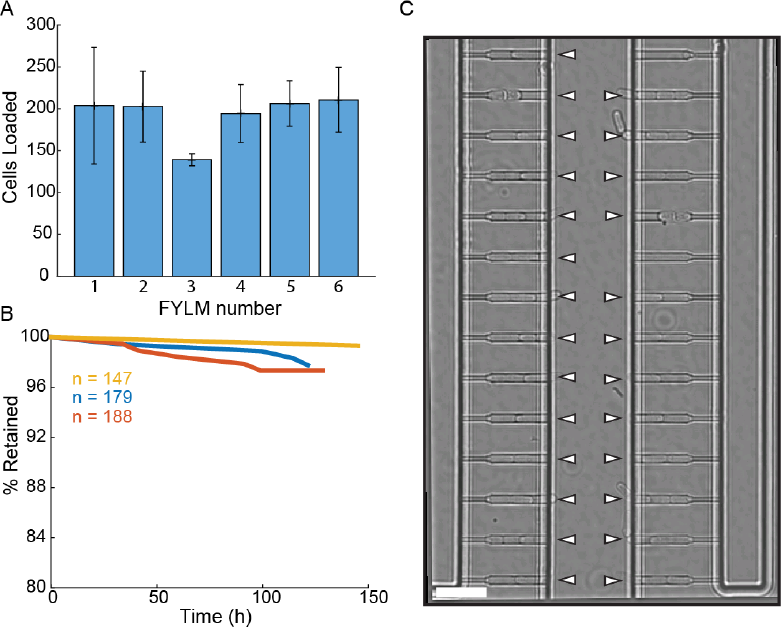
Loading and retention of cells in the multFYLM. (A) Loading efficiency of each FYLM subunit in the multFYLM. Error bars: S.D. of the mean of three loading experiments. (B) Retention efficiency of cells in the FYLM under constant media flow. Three representative experiments shown, with the associated number of loaded cells. We observed retention efficiencies as high as 99% over 140 hours. (C) Image of a single field of view of one FYLM subunit. White triangles mark captured cells. Scale bar: 25 μm.

**Supplemental Figure 3.**
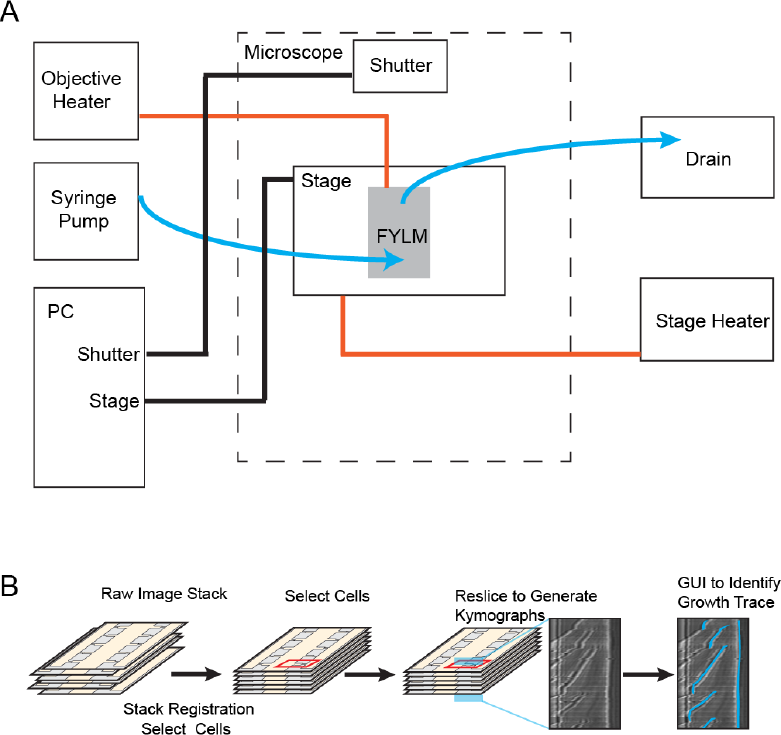
Experimental apparatus and image processing workflow. (A) Schematic of experimental apparatus. Blue lines represent flow of media, red lines represent temperature control, and black lines represent computer-controlled mechanical processes. A syringe pump delivers a constant flow of fresh media to the FYLM. (B) FYLM Critic software workflow. Image stacks are first registered to remove jitter associated with moving the microscope stage. The user then selects cells of interest with a graphical user interface (GUI), and the software generates kymographs. Kymographs are annotated via a semi-automated GUI with the user helping to trace the growth of each cell. The software then quantifies the trace data and exports associated white-light and fluorescence intensities for downstream processing in MATLAB.

**Supplemental Figure 4.**
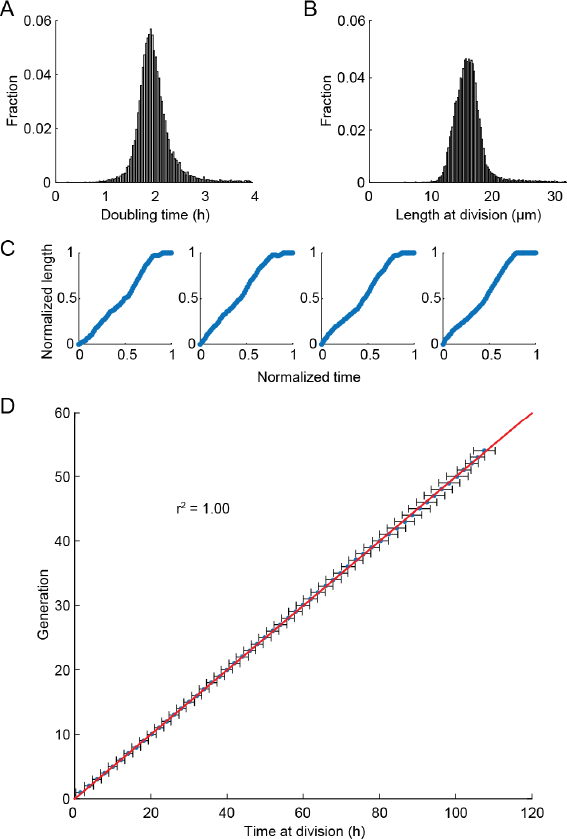
Health of cells in the multFYLM. (A) Histogram of the doubling times and (B) division lengths of wild-type S. pombe (h-972) observed in the FYLM. The doubling time was 2.05 ± 0.45 hours (mean ± S.D., n = 13,453 cells). The mean length at division was 16 ± 2.2 μm. (C) Four representative examples of new end take-off (NETO) in cells growing within the FYLM. Each graph shows the normalized increase in the length of a randomly selected cell over a normalized time comprising one generation. (D) Doubling time did not generally change with age. Horizontal error bars are the SD of the mean time at division for a given generation. Number of cells used ranged from n=517 at generation 1, to n=147 at generation 53. The number of cells declines due mostly to cell death during the experiment.

**Supplemental Figure 5.**
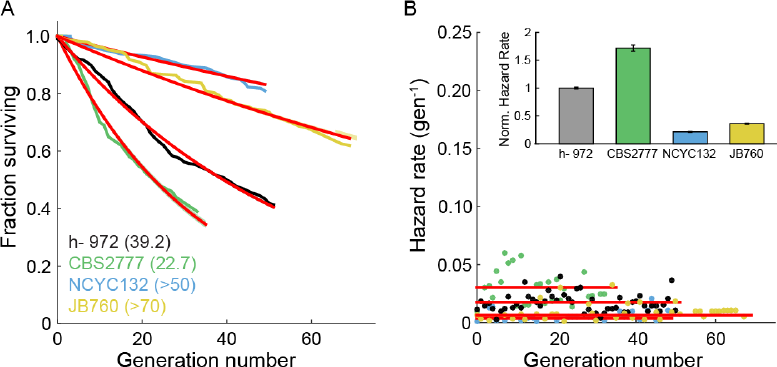
Survival and hazard curves for wild type fission yeast isolates. (A) Survival curves of several wild-type strains evaluated in the FYLM. The replicative half-life is indicated in parenthesis. Red lines: exponential decay fit to the data. For the two particularly long-lived strains NCYC132 and JB760, we could only estimate a lower bound on the mean replicative lifespan. (B) Hazard functions of strains shown in (A). Inset: hazard rates normalized to the wild-type (h-972) hazard rate (error bars indicate 95% CI). The constant hazard rates illustrate that none of the WT populations observed are aging.

**Supplemental Figure 6.**
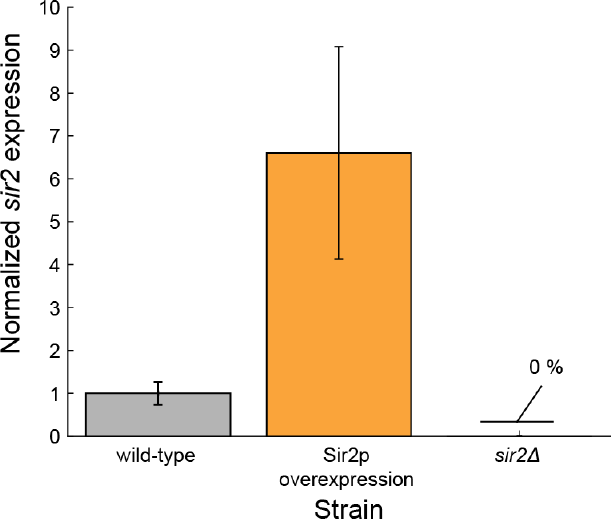
*sir2* expression levels. qPCR results, normalized to wild-type expression. No expression was noted in the *sir2Δ* cells. strain was included as a negative control, no measurable expression was noted. Error bars: normalized SEM of at least 3 replicates.

**Supplemental Figure 7.**
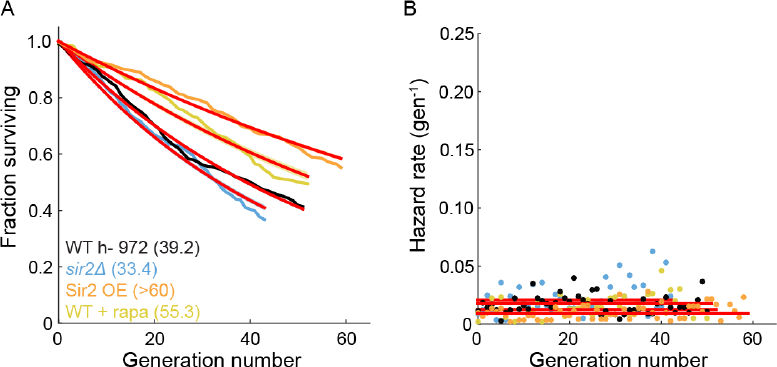
The effect of Sir2p and rapamycin on the RLS of fission yeast. (A) The replicative half-life is indicated in parenthesis. A lower bound is reported for the long-lived Sir2p overexpression strain. Red lines represent an exponential decay fit to the curve. (B) Hazard functions of strains shown in (A). The constant hazard rates illustrate that none of the WT strains are aging.

**Supplemental Figure 8.**
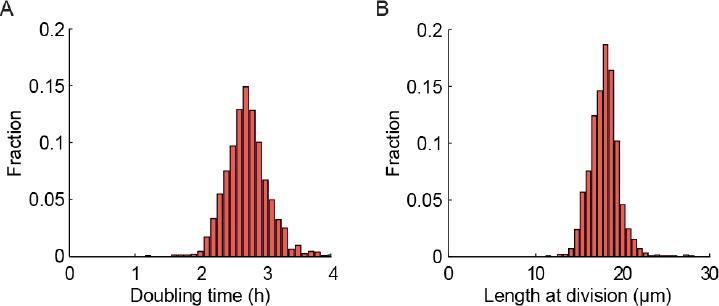
(A) Histogram of doubling times of a strain expressing gar2-mCherry. The doubling time is 2.83 ± 0.87 hours (mean ± st. dev.; n = 1547). (B) Histogram of length at division. The length is 18.0 ± 2.3 μm (mean ± st. dev.; n = 1547).

**Supplemental Figure 9.**
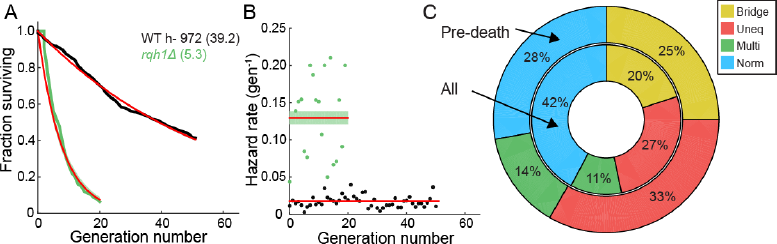
Characterization of the RLS of an *rqh1Δ* strain. (A) Survival curve of WT (black, 95% CI: 38.6-39.8 generations) and *rqh1Δ* strain (green, 95% CI: 5.0-5.7) (B) Hazard curve of WT (black) and rqh1 deletion strain (green). (C) The incidence of nucleolar defects for all divisions (center ring) and preceding death (outer ring).

**Supplemental Table 1.** Comparison of cell metrics to literature. All cells grown in the FYLM were grown with YES media at 31 °C. “Rate Change Point” (RCP) is the position of the new end takeoff rate change in a fractional measurement of cell cycle. “% Change of Rate at RCP” is the change in slope of a linear regression performed on smoothed data after the RCP with respect to the slope of a linear regression performed on smoothed data before the RCP. Measurements are presented as mean ± standard deviation. Cells for reference (1) in the table were grown on an agar substrate made with conditioned EMM3 at 35 °C. Cells for reference (2) in the table were grown using YE4S medium in a microfluidic device at 32 °C. Cells for reference (3) in the table were grown on an agar substrate made with YES (1% Yeast Extract, 3% glucose) at 30 °C

| Measurement | This Research | Literature | Ref |
| --- | --- | --- | --- |
| Length at Birth ($\mu\text{m}$) | $8.0 \pm 1.5$ ($n = 13,905$) | $8.2 \pm 0.52$ ($n = 164$) $8.7 \pm 0.9$ ($n \leq 900$) | (1)<br>(2) |
| Length at Division ($\mu\text{m}$) | $16.0 \pm 2.2$ ($n = 13,456$) | $14.4 \pm 0.85$ ($n = 164$)<br>$14.7 \pm 1.2$ ($n \leq 900$) | (1)<br>(2) |
| Cycle Time (m) | $122 \pm 28$ ($n = 13,456$) | $148 \pm 16$ ($n = 164$) $133 \pm 16$ ($n = 72$) | (1)<br>(3) |
| Rate Change Point | $0.36 \pm 0.07$ ($n = 50$) | $0.34$ ($n = 55$) | (1) |
| % Change of Rate at RCP | $0.22 \pm 0.19$ ($n = 33$) | $0.31$ ($n = 55$) | (1) |

**Supplemental Table 2.** Mean RLS and hazard rates for key strains. Survival curves were fitted with either exponential decay or Gompertz functions, yielding fit values and β ± 95% confidence interval. RLS was calculated using either Equations 6 or 7, range is 95% confidence interval. “n” denotes the number of individual cells used in the experiment. “r^2^_adj_” is the r^2^ value of the fit adjusted for the number of coefficients. NA indicates “not any”

| strain | n | Mean RLS (95% CI) | Age-independent rate (gen<sup>-1</sup>) | Age-dependent rate (gen<sup>-1</sup>) | r<sup>2</sup><sub>adj</sub> |
| --- | --- | --- | --- | --- | --- |
| Wild-type (h- 972) | 440 | 39.2 (38.6-39.8) | 0.017 ± 0.001 | N.A. | 0.99 |
| *rqh1Δ* | 126 | 5.3 (5.0-5.7) | 0.130 ± 0.009 | N.A. | 0.95 |
| WT + rapamycin | 184 | 55.3 (53.1-57.6) | 0.013 ± 0.001 | N.A. | 0.96 |
| *sir2Δ* | 329 | 33.4 (32.6-34.3) | 0.021 ± 0.001 | N.A. | 0.98 |
| *sir2* OE | 301 | > 60 | 0.009 ± 0.001 | N.A. | 0.98 |
| WT (CBS2777) | 100 | 22.7 (21.9-23.6) | 0.031 ± 0.002 | N.A. | 0.98 |
| WT (NCYC132) | 226 | > 50 | 0.003 ± 0.001 | N.A. | 0.97 |
| WT (JB760) | 147 | >70 | 0.006 ± 0.001 | N.A. | 0.98 |
| *S. cerevisiae* | 458 | 25.6 (23.6-28.0) | 0.005 ± 0.001 | 0.106 ± 0.005 | 0.99 |

**Supplemental Table 3.**
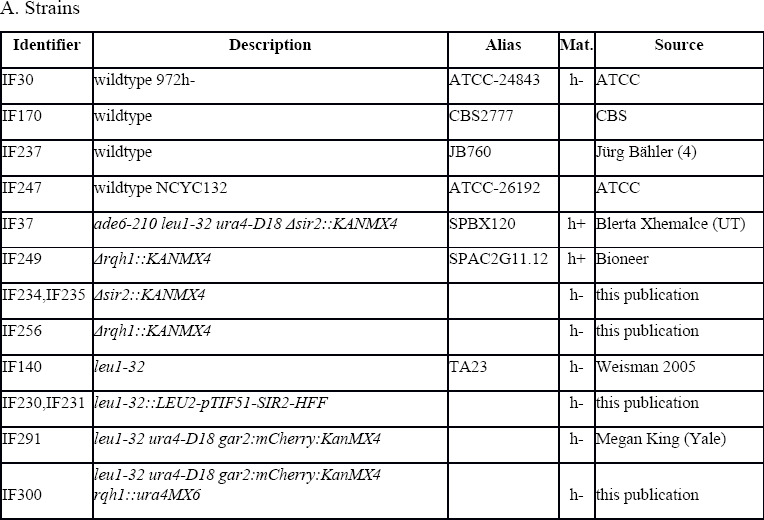
Mean RLS and hazard rates for key strains. Survival curves were fitted with either exponential decay or Gompertz functions, yielding fitvaluesand α ± 95% confidence interval. RLS was calculated using either Equations 6 or 7, range is 95% confidence interval. “n” denotes the number of individual cells used in the experiment. “r^2^_adj_” is the r^2^ value of the fit adjusted for the number of coefficients. NA indicates “not any”

## Supplemental Movie 1. Operation of the multFYLM.

Time-lapse imaging of fission yeast in a single field of view of the FYLM for approximately 140 hours. Scale bar is 20 μm.

## Supplemental Movies 2-4. New-pole sibling phenotypes.

Movie S2 is of a cell where the last division produces a healthy new-pole cell that divides more than once, Movie S3 is of a cell where the last division produces an unhealthy new-pole cell that divides once before dying, Movie S4 is of a cell where the last new-pole cell does not divide. Scale bars are 20 μm.

